# Restoration of Type 17 immune signaling is not sufficient for protection during influenza-associated pulmonary aspergillosis

**DOI:** 10.1101/2024.07.01.601559

**Authors:** Aijaz Ahmad, Ravineel Bhan Singh, Kara Nickolich, Matthew Pilewski, Caden Ngeow, Kwame Frempong-Manso, Keven Robinson

## Abstract

Influenza-associated pulmonary aspergillosis (IAPA) is a severe complication of influenza infection that occurs in critically ill patients and results in higher mortality compared to influenza infection alone. Interleukin-17 (IL-17) and the Type 17 immune signaling pathway cytokine family are recognized for their pivotal role in fostering protective immunity against various pathogens. In this study, we investigate the role of IL-17 and Type 17 immune signaling components during IAPA. Wild-type mice were challenged with influenza A H1N1 (Flu) and then exposed to *Aspergillus fumigatus* ATCC42202 resting conidia on day 6 post-influenza infection, followed by the quantification of cytokines and chemokines at 48 hours post-fungal infection. Gene and protein expression levels revealed that IL-17 and Type 17 immune cytokines and antimicrobial peptides are downregulated during IAPA compared to mice singularly infected solely with *A. fumigatus*. Restoration of Type 17 immunity was not sufficient to provide protection against the increased fungal burden observed during IAPA. These findings contrast those observed during post-influenza bacterial super-infection, in which restoration of Type 17 immune signaling protects against exacerbation seen during super-infection. Our study highlights the need for future studies to understand the immune mechanisms that increase susceptibility to fungal infection.

**Importance:** IAPA significantly elevates the risk of mortality in patients with severe influenza. Type-17 immunity is critical to host defense during fungal infections and, therefore, vital to understand its role during IAPA. The observations in this study reveal that Type 17 immunity is impaired during IAPA, potentially increasing susceptibility to secondary infection with *Aspergillus fumigatus*. However, restoration of IL-17 signaling alone is not sufficient to reduce fungal burden in our murine IAPA model. These observations differ from those observed in post-influenza bacterial super-infections, suggesting that the mechanisms underlying viral-fungal super-infection are different than those that underly viral-bacterial super-infection. By elucidating the complex interactions between the host immune system, influenza, and *A. fumigatus*, these findings are vital for developing strategies to enhance immune responses and improve survival rates during IAPA.

## INTRODUCTION

Influenza-associated pulmonary aspergillosis (IAPA) is a severe complication of influenza infection that occurs in critically ill patients and results in higher mortality compared to influenza infection alone [1]. The pathology of IAPA manifests through invasive growth of *Aspergillus fumigatus* within the lungs during influenza infection [2]. Influenza virus damages the respiratory epithelium, compromising the barrier function and allowing opportunistic fungi such as *A. fumigatus* to invade and colonize the lung tissue. The dysregulated immune state of the host, caused by the viral infection, provides an optimal environment for fungal growth and dissemination.

The interleukin-17 (IL-17) cytokine family is a pivotal component in mediating protective immunity against various pathogens [3]. IL-17, primarily produced by a subset of T helper cells known as Th17 cells, has been implicated in the immune response against extracellular pathogens. IL-17 exerts its immunomodulatory effects by promoting the synthesis of pro-inflammatory molecules, including cytokines, chemokines, and antimicrobial peptides. These molecules collectively facilitate the recruitment and activation of neutrophils and other immune effectors to sites of infection, thereby augmenting the host defense against invading pathogens [4].

Despite the well-established role of IL-17 in immunity against various pathogens, its specific involvement in the context of IAPA remains understudied. Therefore, understanding the immune response in this context is vital for improving therapeutic strategies and patient outcomes. Our current study investigates the role of IL-17 and IL-17-related cytokines in IAPA, employing a murine model to delineate their impact on disease pathogenesis and immune responses. Our observation sheds light on the complex host immune system that occurs during IAPA, aiming to uncover novel therapeutic avenues and enhance patient outcomes in managing IAPA.

## MATERIALS AND METHODS

### Animals

Six to eight-week-old male C57BL/6 mice were purchased from Taconic Farms (Germantown, NY). The mice were kept in a pathogen-free environment and co-housed in the same facility before the commencement of the studies. All animal studies were performed according to the protocol for the care and use of animals sanctioned by the University of Pittsburgh Institutional Animal Care and Use Committee. All the studies used age- and sex-matched mice.

### Pathogens and superinfection model

Influenza A/PR/8/34 H1N1 was propagated in chicken eggs as previously described [5] or by using Madin-Darby canine kidney (MDCK) cells. The cells were maintained in DMEM with 10% FBS (Bio-Techne, Minneapolis, MN), penicillin (100 U/ml), streptomycin (100 ug/ml) (Invitrogen, Waltham, MA). The cells were washed with PBS, and infected 0.001 MOI of influenza virus A/Puerto Rico/8/1934 (H1N1) in DMEM with 0.2% bovine serum albumin (Invitrogen, Waltham, MA), and 2 μg/ml of L-tosylamido-2-phenyl ethyl chloromethyl ketone (TPCK) (Sigma-Aldrich, MO). The virus containing supernatant was harvested after 72 hours and the viral titer was determined by standard plaque assay. Mice were infected with 100 PFU of influenza A/PR/8/34 H1N1 (in 50 μl sterile PBS) from a frozen stock or control PBS by oropharyngeal aspiration. Infected mice were incubated for 6 days, at which time mice received 2.5 × 10^7^ conidia of *A. fumigatus* ATCC42202 inoculum or PBS control. At 48 hours post-fungal infection, all the mice were euthanized to harvest lungs for further studies.

### Lung inflammation analysis

After harvesting, mouse lungs were lavaged with 1 ml of sterile PBS to perform inflammatory cell differential counts. The upper lobe of the right lung was homogenized in sterile PBS for counting fungal colonies and cytokine analysis, conducted either with Lincoplex (Millipore, MO, USA) or ELISA assays (R&D Systems, MN, USA), following the manufacturer’s guidelines. The middle and lower lobes of the right lung were snap-frozen and then homogenized under liquid nitrogen for RNA extraction using the RNA isolation Kit (Agilent Technologies, TX, USA). The RNA analysis was carried out *via* standard RT-qPCR employing Bio-Rad SSO advanced Universal Probes Supermix (CA, USA). Gene expression analysis was performed from two replicate samples. It was calculated using the formula ΔCq=2^Cq target gene-Cq reference gene^, where the quantitation cycle (Cq) was the average Cq value of the target gene minus the *HPRT* reference gene’s mean.

### Flow cytometry

Flow cytometry analysis was conducted on the whole left lung. After harvesting, the left lung underwent collagenase digestion, following a previously described protocol [6]. The resulting single-cell preparations were *in vitro* stimulated with PMA (50 ng/ml) and ionomycin (750 ng/ml) for four h at 37 °C. Subsequently, cells were stained with antibodies, fixed and permeabilized, and stained with fluorescent-conjugated antibodies (BD Biosciences). The analysis used a Cytek Aurora™ CS System (Cytek® Biosciences Bethesda, MD, USA).

## RESULTS

### Type 17 cytokines are inhibited during IAPA

We have previously published a murine model of IAPA that demonstrates increased morbidity in mice co-infected with influenza and *A. fumigatus* [7]. Type 17 immunity plays a critical role in host defense against *A. fumigatus* and other fungal pathogens [8-12]. IL-17 and other Type 17 immune cytokines also play a critical role in the development of bacterial super-infection during influenza [13-14]. Therefore, we hypothesized that the Type 17 immune response would play an essential role during IAPA. To model this hypothesis, C57BL/6J male mice were challenged with a sublethal dose of influenza A PR/8/34 H1N1 (100 PFU) for 6 days, followed by 2.5 × 10^7^[*A. fumigatus* (ATCC strain 42202) conidia, and after 48 hours, fungal burden was assessed. Mice super-infected with influenza and *A. fumigatus* had decreased expression of IL-17, IL-22, IL-23, and IL-1β compared to those infected with *A. fumigatus* alone (**Fig 1A**). This downregulation could potentially contribute to increased morbidity, mortality, and fungal burden compared to singular infection with *A. fumigatus* alone. In addition to the gene expression changes, protein expression analysis also showed a decreased production of IL-17, IL-22, IL-23, and IL-1β proteins in super-infected mice (**Fig 1B)**.

**Figure 1:**
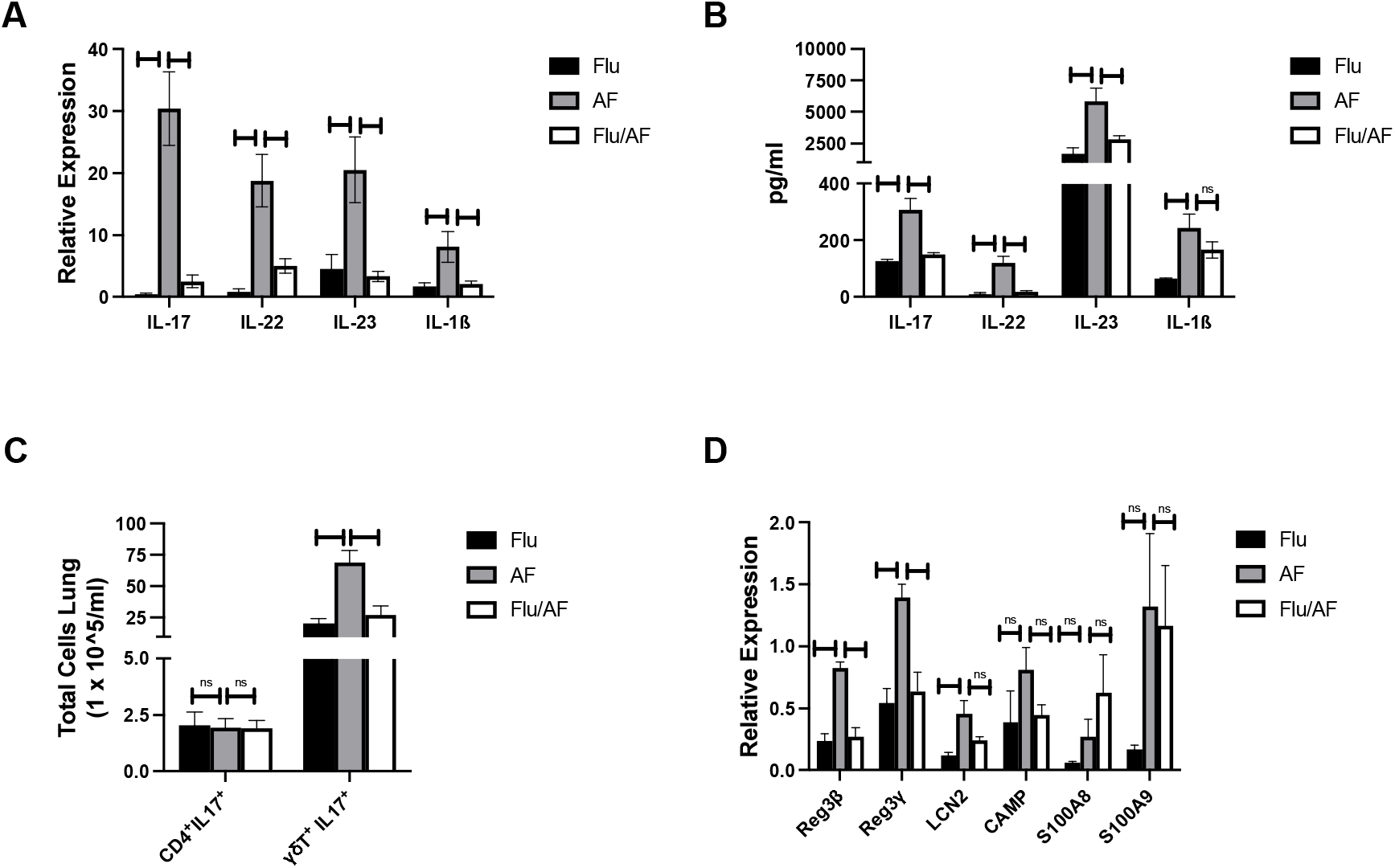
Type 17 immune pathway is downregulated during influenza-associated pulmonary aspergillosis (IAPA). WT mice were infected with influenza A H1N1 PR/8/34 on day 0 and subsequently challenged with 2.5 × 10^7^ *aspergillus fumigatus* (AF) at 6 dpii. Lung samples were collected at 48 h post-AF challenge. (**A**) IL-17 and related pro-inflammatory cytokine gene expression quantified by RT-qPCR, (**B**) Protein levels of IL-17-related cytokines quantified using Lincoplex assay or ELISA, (**C**) Flow cytometry analysis of IL-17-producing cells, CD4^+^ IL-17^+^ T cells and γδ T cells, (**D**) IL-17/IL-22-associated antimicrobial peptide gene expression quantified by RT-qPCR. Data were compiled from two independent experiments and are presented as means ± SEM, with statistical significance marked as \**p* < 0.05, ** *p* < 0.005, and *** *p* < 0.0005 by unpaired student’s T-test.

### Decreased IL-17-producing γδ T cells during IAPA

Flow cytometry analysis was conducted to quantify IL-17-producing T cells in our model. We measured IL17^+^ CD4^+^ T and γδ T cells in our model due to their well-established role as primary producers of IL-17. We observed a notable decrease in the total number of IL-17 producing γδ T cells in super-infected mice compared to singular *A. fumigatus* infection (**Fig 1C**). Comparatively, we observed no difference in the abundance of IL17^+^ CD4^+^ T cells between the different groups (**Fig 1C**). When interpreting these results, it is important to consider the timing of mouse harvesting post-*A. fumigatus* infection. Specifically, the analysis was conducted at a 48-hour time point post-infection. At this early stage, the immune response is still evolving, with distinct subsets of immune cells playing varying roles. In particular, it is noted that there were more innate γδ T cells present at this time point, with T cells potentially infiltrating at later stages of infection.

### Reduced expression of IL-17/IL-22 associated antimicrobial peptides in IAPA

As IL-17 and IL-22 are distinct lineages of Type 17 cells [15], we studied their effector function by examining gene expression levels of IL-17- and IL-22-associated antimicrobial peptides in our model. The expression levels of *Reg3*β, *Reg3*γ, *and Lcn2* were significantly reduced in super-infected mice compared to those infected solely with *A. fumigatus* (**Fig 1D)**. This reduction of IL-17- and IL-22-associated antimicrobial peptides may play a role in fungal clearance during IAPA.

### Restoration of Type 17 immunity does not enhance fungal clearance during IAPA

With the observation that Type 17 cytokines and antimicrobial peptides were significantly downregulated during IAPA compared to *A. fumigatus* alone, we hypothesized that their restoration could provide protection during IAPA. To test this hypothesis, we overexpressed IL-17 proposing that it would rescue Type-17 immunity and enhance fungal clearance. Overexpression of IL-17 significantly upregulated the levels of IL-17a mRNA (**Fig 2B**) and downstream inflammatory mediators TNF⍰ and CXCL1 during IAPA (**Fig 2C**) [16,17]; however, fungal burden remained unchanged **(Fig 2A)**. Interestingly, the total number of inflammatory cells in the bronchoalveolar lavage fluid and the total numbers of neutrophils and macrophages measured by cytospin differential also remained unchanged with upregulation of Type 17 immunity (**Fig S1**). Additionally, we administered murine recombinant IL-17 protein during IAPA and fungal burden was unchanged (**Fig S2**). Next, we restored components of the Type 17 immune signaling pathway that are both upstream and downstream of IL-17 to determine effects on fungal clearance. Since both IL-1β and IL-23 are known to induce IL-17 production from γδ T cell [18], and we observed decreased production during IAPA (**Fig 1A-B**), we administered IL-1β and IL-23+IL-1β together for synergistic effects during IAPA. There was a significant increase of IL-17 mRNA in the mice that received exogenous IL-1β and IL-23/IL-1β (**Fig 2E**); however, fungal burden remained unchanged (**Fig 2D**). Additionally, IL-1β and IL-23+IL-1β did not alter IL-22 mRNA expression (**Fig 2E**). Super-infected mice were administered either with Reg3β or Reg3γ to explore potential therapeutic interventions. Although both these peptide levels have been reduced in super-infected mice, neither changed fungal clearance during IAPA (**Fig 2F**). Additionally, we observed no difference in the total cell count of immune cells in bronchoalveolar lavage fluid (**Fig S3)**. Collectively, these results indicate that restoration of IL-17 signaling is not sufficient to restore fungal clearance during IAPA. These findings suggest the involvement of mechanisms beyond Type 17 inhibition causing decreased fungal clearance during IAPA and highlight the complexity of IAPA pathogenesis.

**Figure 2:**
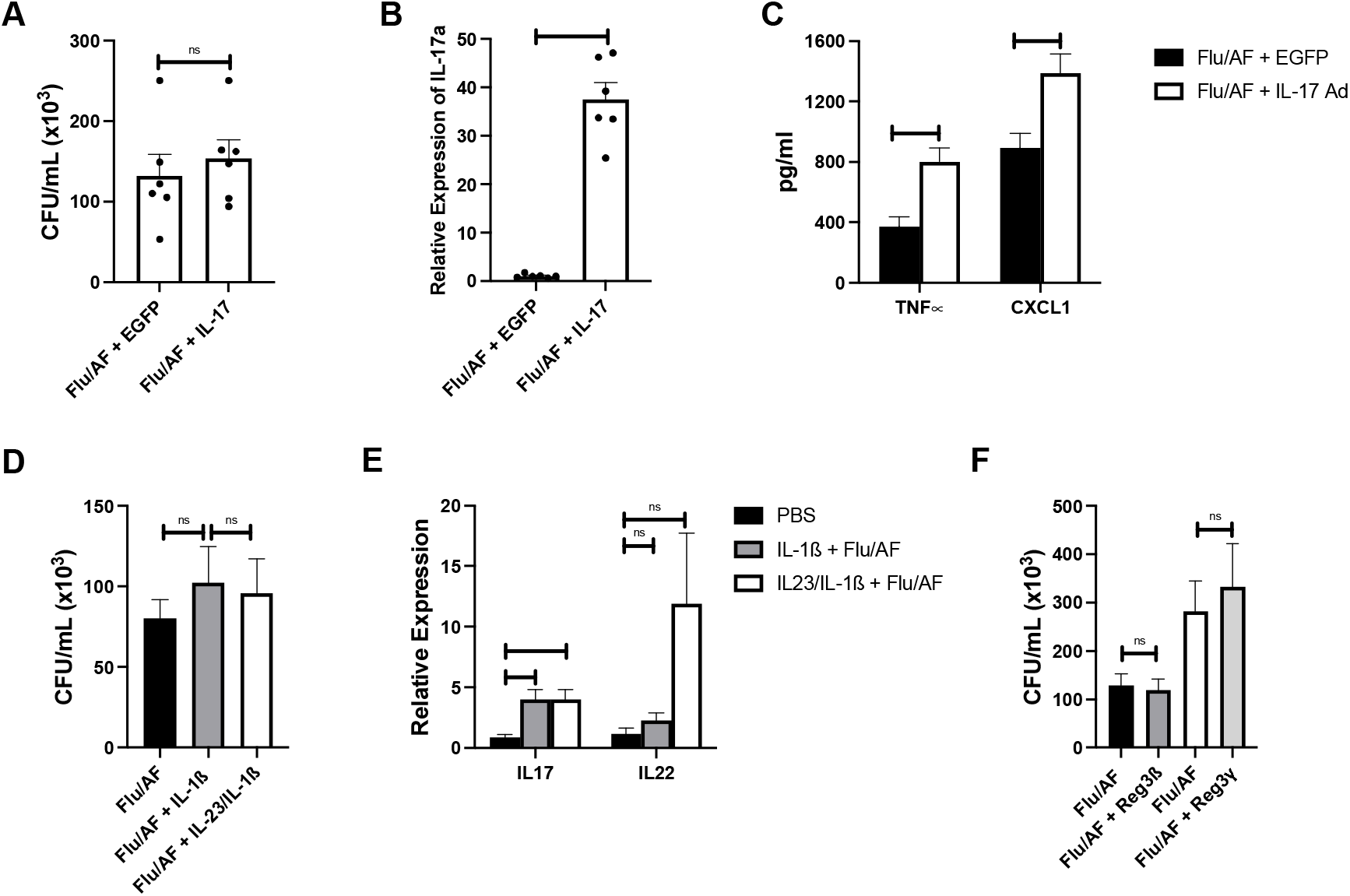
Restoration of Type 17 immune signaling is not sufficient to provide protection during IAPA. WT mice were infected with influenza A H1N1 PR/8/34 on day 0 and subsequently challenged with 2.5 × 10^7^ *aspergillus fumigatus* (AF) at 6 dpii. Lung samples were collected at 48 h post-AF challenge. (**A-C**) Mice were administered IL-17 expressing adenovirus (IL-17) or control adenovirus during IAPA and (**A**) fungal burden measured by CFU, (**B**) IL-17a gene expression quantified by RT-qPCR, and (**C**) TNF_IZI_ and CXCL1 cytokines measured by ELISA. (**D-E**) Mice were administered IL-1β or IL-1β+IL-23 murine recombinant protein during IAPA and **(D)** fungal burden measured by CFU, (**E**) IL-17a and IL-22 gene expression quantified by RT-qPCR. **(F)** Mice were administered Reg3β or Reg3γ murine recombinant proteins during IAPA and fungal burden measured by CFU. Data were compiled from two independent experiments and are presented as means ± SEM, with statistical significance marked as \**p* < 0.05, ** *p* < 0.005, and **** *p* < 0.0001 by unpaired student’s T-test.

## DISCUSSION

Although viral-fungal co-infections are associated with high mortality, limited data exists regarding pathophysiology and lung immunology. Influenza and *A. fumigatus* have individually been studied due to their clinical significance; however, the synergy and complexities that arise when these pathogens co-exist within the same host remain poorly understood. Invasive pulmonary aspergillosis was classically considered a disease of immunocompromised patients; however, recent clinical observations have reported invasive pulmonary aspergillosis following influenza infection (IAPA) in immunocompetent patients [19]. The increased risk of developing aspergillosis in patients with preceding influenza can be partially attributed to viral-induced epithelial disruption, the first line of host defense against fungal infections. However, recent evidence suggests that second (phagocytosis and the killing of *Aspergillus* conidia by phagocytes) and third lines (extracellular mechanisms, mediated by neutrophils, to kill the *Aspergillus*) of the antifungal host responses are also impaired in patients with IAPA [1,20]. IAPA has also been documented to provoke a severe inflammatory response, resulting in a cytokine storm within lung tissue [21]. IAPA has been described for decades and has been increasingly recognized since the 2009 H1N1 influenza pandemic. Notably, *Aspergillus* species are also implicated in causing super-infection during SARS-CoV-2 (COVID-19) infections [22,23], underscoring the importance of studying viral-fungal super-infections. Additionally, studies suggest that the hyperinflammatory responses driven by systemic cytokines in COVID-19 patients also contribute to CAPA [24].

The Type 17 cytokine family is recognized for its pivotal role in fostering protective immunity against a spectrum of pathogens [25]. Previous research has substantiated the involvement of IL-17 in the context of viral-bacterial superinfections [13]. The IL-17 pathway has also promoted *Aspergillus* clearance within pulmonary tissues [8-10]. Mechanistically, the protective role for IL-17 is mediated by the recruitment of neutrophils through chemokine signaling and the production of antimicrobial peptides (AMP) production [26]. This study aims to delineate the specific roles of IL-17 and Type 17 immune-associated cytokines and chemokines in IAPA.

Our results demonstrate that preceding influenza infection impairs Type17 immunity during IAPA. Both gene expression and protein quantification of key Type 17 immune cytokines, particularly IL-17 and IL-22, are decreased during IAPA compared to singular infection. IL-17, a pro-inflammatory cytokine, plays an essential role in fungal infections by recruiting neutrophils and other immune cells to the site of infection and by inducing the production of antimicrobial peptides [27]. Additionally, IL-17 synergistically collaborates with IL-22 for a robust immune response against fungal infection [28]. The synergistic action of IL-17 and IL-22 is crucial for a robust immune response, and their inhibition can lead to a compromised ability to control fungal infections. Decreased IL-17 production was also seen by Lee et al., using a similar murine model of IAPA; however, we also show a reduction in other Type 17-immune associated cytokines and antimicrobial peptides [29]. IL-1β, a pro-inflammatory cytokine critical to Type 17 immunity, is reduced in super-infection compared to singular *A. fumigatus* challenge. IL-1β has also been reported to induce neutrophils and macrophages recruitment to lungs during microbial invasion [7] and stimulates endothelial adhesion molecules, different cytokines and chemokines, and the Th17 adaptive immune response [30]. Our results align with the previous findings of downregulation of *IL1B* that was observed in humans during IAPA [20]. Furthermore, in our murine model of IAPA, the impaired production of IL-17 and IL-22 is also associated with reduced levels of antimicrobial peptides, weakening the host’s defense mechanisms. These findings highlight that the downregulation of Type 17 immunity during IAPA compromises the host’s defense mechanisms and thereby increasing susceptibility to *Aspergillus* infections.

Despite the observed reduction in Type 17 immunity during IAPA, restoration of various components of the IL-17 signaling pathway, both upstream and downstream of IL-17, did not lead to improved fungal clearance. In contrast to prior studies that showed restoration of Type 17 immune pathway components rescued bacterial clearance during post-influenza bacterial super-infection [13,14,31], the current study indicates that restoration of IL-17 signaling alone is not sufficient to reduce fungal burden in a murine IAPA model. IL-17 may play a protective role but not a restorative role during IAPA. Notably, the immune response is dynamic during infection and augmentation of Type 17 signaling at other time points during IAPA may produce different results. Interestingly, downregulation of IL-1β in our mouse model is consistent with the findings in human patients with IAPA versus influenza alone [20]. This consistency between our findings in mice and human patients strengthens the validity of our results and suggests that the observed decrease in cytokines is a robust effect that is relevant across species. Importantly, these studies show that immune regulation during post-influenza fungal super-infection and post-influenza bacterial super-infection are not driven by the same mechanisms. It underscores the need for additional studies to understand the immune mechanisms that increase susceptibility to fungal infection during influenza and how delineation of the specific cell types and immune pathways that are necessary for fungal host defense during viral infection may lead to future therapeutics.

## ETHICS APPROVAL

The University of Pittsburgh Institutional Animal Care and Use Committee approved the study under Protocol No. 21063690.

## ACKNOWLEDGMENTS

The authors appreciate the National Institute of Allergy and Infectious Diseases Grant R01 (Award No. R01AI153337 to KMR) and the National Heart, Lung and Blood Institute Grant K08 (Award No. K08HL133445 Award No to KMR).

## DATA AVAILABILITY

All the associated data is provided with this manuscript or in supplementary files.

## SUPPLEMENTAL MATERIAL

Supplemental Figures S1-S3.

