## Supplemental Figs for "Restoration of Type 17 immune signaling is not sufficient for protection during influenza-associated pulmonary aspergillosis"

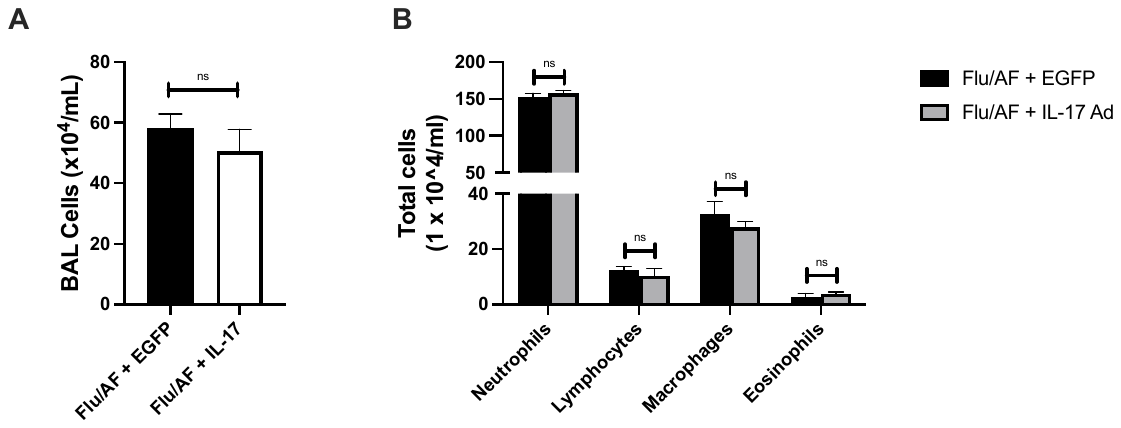

**Figure S1**: Bronchoalveolar lavage (BAL) cell counts (**A**) and BAL differentials (**B**) were measured in wild-type mice infected with influenza A/PR/8/34 H1N1 (Flu) and *A. fumigatus* ATCC42202 resting conidia (AF). The administration of exogenous IL-17 adenovirus (IL-17) did not significantly affect BAL cell counts or any of the differential cell populations tested. Statistical significance was assessed using an unpaired Student’s t-test. Each experiment was independently performed twice, and the data presented are combined from these experiments.

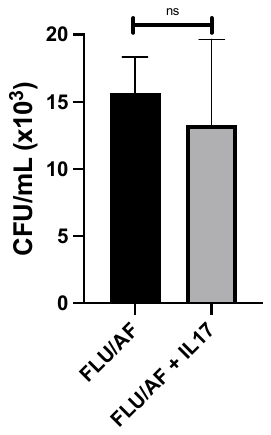

**Figure S2**: Fungal load in the lungs was determined by CFU plating in wild type mice infected with Flu and AF. The administration of recombinant IL-17 protein did not significantly rescue the co-infected mice, as indicated by the lack of significant differences in CFU counts across the different treatment groups. The experiment was conducted twice, and the data are presented as means ± SEM.

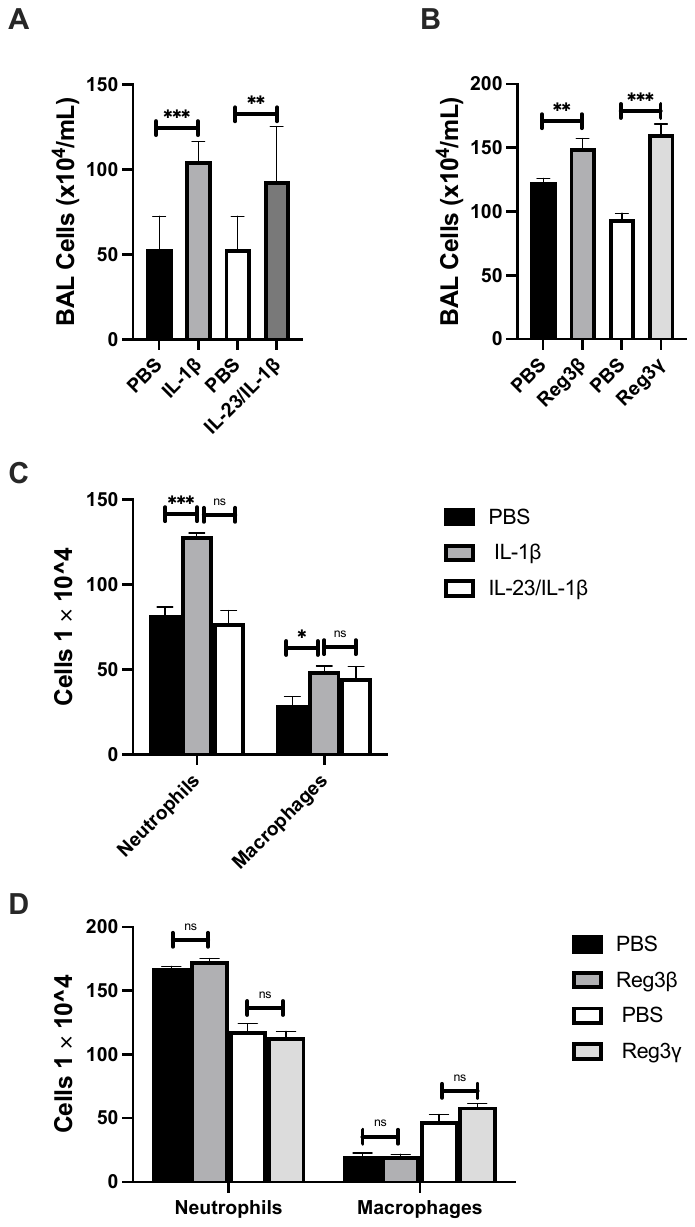

**Figure S3**: Measurement of Bronchoalveolar lavage (BAL) cell counts in absence (PBS control) and presence of (**A**) different cytokines (IL-1β and IL-23/IL-1β) and (**B**) antimicrobial peptides (Reg3β or Reg3γ) in wild-type mice infected with influenza A/PR/8/34 H1N1 (Flu) and *A. fumigatus* ATCC42202 resting conidia (AF). The administration of IL-1β and IL-23/IL-1β, but not the antimicrobial peptides, resulted in a significant increase in BAL cell counts in the context of IAPA. In addition, the impact on differential cell populations was recorded in presence of (**C**) IL-1β and IL-23/IL-1β and (**D**) Reg3β or Reg3γ. None of the externally provided factors except L-1β significantly affect the differential cell populations tested. This experiment was independently repeated twice, with data presented as means ± SEM. Statistical significance was determined using an unpaired Student’s t-test, with ***p* < 0.005 and ****p* < 0.0005.
